# Mechanical properties of breast, kidney, and thyroid tumours measured by AFM: relationship with tissue structure

**DOI:** 10.1101/2022.06.09.495321

**Authors:** A. Levillain, C.B. Confavreux, M. Decaussin-Petrucci, E. Durieux, P. Paparel, K. Le-Bail Carval, L. Maillard, F. Bermond, D. Mitton, H. Follet

**Author notes:** Corresponding authors: David Mitton, Laboratoire de Biomécanique et Mécanique des Chocs UMR_T 9406 (Univ Eiffel-Univ Lyon 1), Université Gustave Eiffel, Campus de Lyon, 25 Avenue François Mitterrand, 69675 BRON Cedex FRANCE, Hélène Follet, INSERM, Université de Lyon, UMR1033, Faculte de Medecine Lyon Est-Domaine Laennec-6th, 7-11, rue G. Paradin, 69372 Lyon cedex 08, France.

## Abstract

The mechanical properties of the extracellular matrix are essential for regulating cancer cell behaviour, but how they change depending on tumour type remains unclear. The aim of the current study was to determine how the mechanical properties of tumours that frequently metastasize to bones were affected depending on histological type. Human breast, kidney, and thyroid specimens containing tumour and normal tissue were collected during surgery. The elastic modulus and elastic fraction of each sample were characterised using atomic force microscopy and compared with histopathological markers. We observed that tumour mechanical properties were differentially affected depending on organ and histological type. Indeed, clear cell renal carcinoma and poorly differentiated thyroid carcinoma displayed a decrease in the elastic modulus compared to their normal counterpart, while breast tumours, papillary renal carcinoma and fibrotic thyroid tumours displayed an increase in the elastic modulus. Elastic fraction decreased only for thyroid tumour tissue, indicating an increase in the viscosity. These findings suggest a unique mechanical profile associated with each subtype of cancer. Therefore, viscosity could be a discriminator between tumour and normal thyroid tissue, while elasticity could be a discriminator between the subtypes of breast, kidney and thyroid cancers.

**Graphical abstract:** 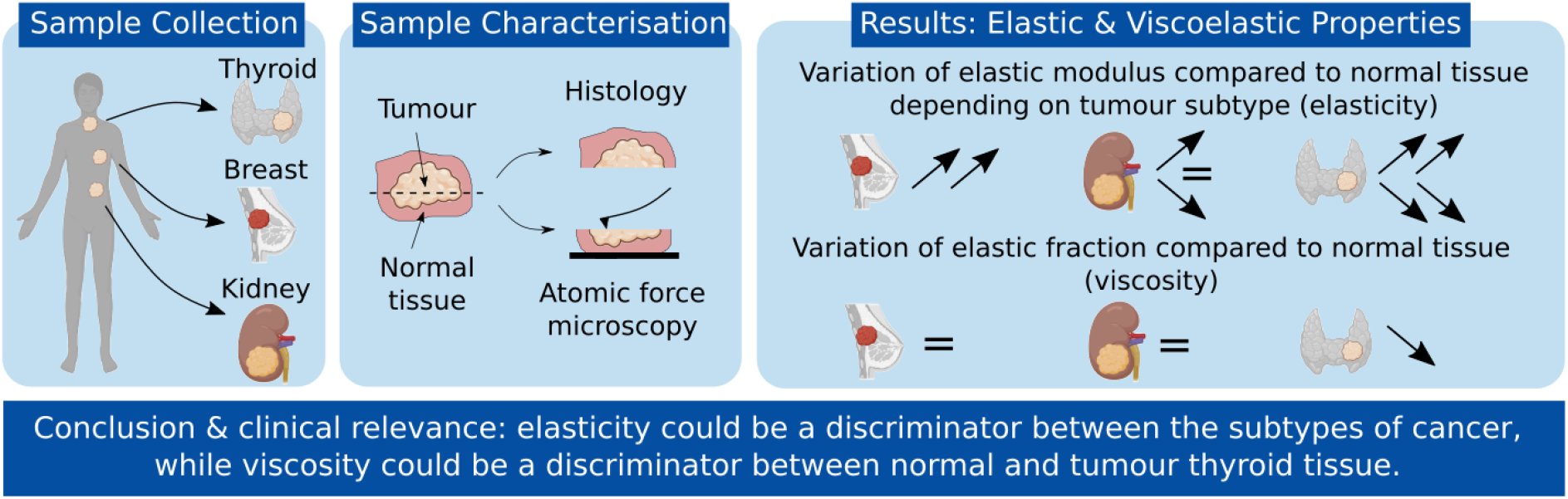

## 1 Introduction

Bone metastases occur in more than 1.5 million cancer patients worldwide, and are associated with serious skeletal-related events, including pathological fracture, pain, disability, and spinal cord compression (*1, 2*). The most common cancers to metastasize to bone are breast, prostate, thyroid, lung, and kidney cancers (*3*). The ability of cancer cells to metastasize mainly depends on the physical interactions and mechanical forces between these cells and the microenvironment they migrate through. In particular, the extracellular matrix (ECM) plays a crucial role in the metastatic spread. The ECM has unique physical, biochemical, and biomechanical properties that are essential for regulating cell behaviour. During cancer progression, the ECM undergoes extensive remodelling (*4*) and its stiffness plays a causative role in cancer pathogenesis (*5*). In breast, a stiffer ECM stimulates epithelial-like cell transformation from normal cells to malignant cells with a more aggressive phenotype that promotes cancer cell invasion (*4, 6*). However, although several studies have been conducted on the mechanical characterisation of breast tumour at the tissue level (*7–10*), there is lack of knowledge on the mechanical properties of tumours from other tissues, including thyroid and kidney.

Several lines of evidence suggest that the extent of mechanical alterations depends on the type of tumour. First, while most studies on breast cancer have shown that tumour tissue exhibited an increase in the stiffness compared to normal tissue (*7, 9, 11, 12*), a study using a palpation device to measure tissue elasticity showed that renal cell carcinoma tissue was softer than normal kidney tissue (*13*).Differences in mechanical alterations between tumour and normal tissue have also been observed for a same organ. In their study characterising the mechanical properties of breast tissue samples from different types, Samani *et al* found that the increase in the elastic modulus was dependent on the tumour type, with high-grade invasive ductal carcinoma exhibiting the highest increase compared to normal tissue (*9*). Similarly, the elastic modulus of thyroid tumour tissue is differentially affected depending on the type of cancer, with samples of papillary carcinoma exhibiting a significantly higher elastic modulus compared to normal thyroid tissue, while samples of follicular adenocarcinoma exhibit a similar stiffness compared to normal thyroid tissue (*14*). Second, *in vitro* studies demonstrated that rigidity profile is an intrinsic property of each cancer line; some cells display an increased growth as ECM rigidity increases, while some cancer cells grow equally well across a large spectrum of ECM rigidity (*15*). This finding suggests that cancer spread depends both on the ECM mechanical properties and on the tumour type. It is, therefore, important to characterise the mechanical properties of the tumour along with its histological type. However, a direct comparison from the literature of the mechanical properties of different types of tumours is challenging, because most studies focused on one type of tissue and the techniques of characterisation varied among the studies. More specifically, there is a variability in the length scales between the different techniques of characterisation, from tissue-level (*7, 10*) to tumour-level (*9, 11, 12, 14*). A characterisation of the tumours at the tissue level is more relevant in order to uncover the mechanobiology mechanisms of cancer progression and metastatic spread.

Indentation using atomic force microscopy (AFM) has been used to reveal the nanomechanical signature of human breast (*7, 10*), liver (*16*), brain (*17*), ovarian (*18*), and prostate (*19*) cancer. Mapping of the elastic modulus in the breast tumour tissue at two different stages of cancer progression highlighted different stiffness profiles, with malignant tissues showing a broader distribution of Young’s modulus compared to normal and benign tissues (*7*). This finding emphasizes the ability of AFM to reveal differences in the elastic properties according to cancer progression, and suggests that it could also reveal differences according to tumour type. To date, most research on the mechanical properties of tumour tissue has focused on elasticity as the key determinant of tissue behaviour. Yet, ECM exhibits both elastic and viscous properties, and changes in ECM viscoelasticity affects cell activities (*20*). For example, a decrease in collagen fibers’ stress relaxation is associated with a reduction of cell persistent migration (*21*), highlighting the need to account for both elastic and viscous properties of the tumour. In recent years, significant development has been made for measuring the viscoelastic properties of soft tissues using AFM (*22*). This technique has the potential to reveal the viscoelastic properties of tumour tissue, which can then be correlated with histopathological markers.

The aim of this study was to determine how the mechanical properties of tumour tissues that frequently metastasize to bone are affected depending on their histological type. Based on the findings on breast tumours (*7, 9*), it was first hypothesized that the elastic properties of the tumours are differentially affected depending on their type. Second, given the role of ECM viscoelasticity in regulating cell and tissue dynamics (*20*), it was hypothesized that tumour tissue exhibits a change in viscous properties compared to normal tissue. To test these hypotheses, we characterised the elastic and viscoelastic properties of breast, kidney and thyroid tumour tissues using AFM, and we compare them with histopathological markers.

## 2 Materials and methods

### 2.1 Study participants and tissue preparation

The study protocol has been approved by the French Ethics Committee (CPP SUD-EST 1 France) under the registration number ID-RCB: 2019-A01202-55. All procedures have been conducted in compliance with national and European regulations. All included patients received a clear information and provided a written consent. Human breast (n=6), kidney (n=5), and thyroid (n=4) specimens containing tumour tissue were collected post-surgery at Hospices Civils de Lyon (Groupement Hospitalier Est (KLBC) and Hôpital Lyon Sud (LM, PP), France). From each removed specimen, the pathologist prepared one sample containing healthy and tumour tissue (breast) or one healthy tissue sample and one tumour tissue sample (kidney and thyroid) or for AFM testing. The samples used for AFM were free of necrotic areas and their dimensions were at least 2 x 10 x 10 mm. The remaining specimen was used for routine pathological diagnosis. The samples used for AFM testing were immediately frozen at −80°C and stored at this temperature until the day of experiment.

### 2.2 Histopathological examination

Samples used for pathological diagnosis were fixed in formol and embedded in paraffin. 3 μm-thick sections were cut using a microtome and stained with eosin hematoxylin saffron. For each sample, one section containing tumour tissue was imaged in transmitted illumination using a light microscope. Histopathological examination included assessing the type of carcinoma, extent of fibrosis and necrosis.

### 2.3 AFM testing

Frozen samples were carefully cut using a scalpel to get a 2-3 mm thick slice, with smooth and uniform surfaces. On the day of experiment, samples were thawed at room temperature for 1 hour, glued on a petri dish and immersed in phosphate buffered saline (PBS). Mechanical manipulations during sample preparation were always kept minimal. The samples were then allowed to set for 30 minutes to avoid swelling of the sample during AFM testing. The petri dish was placed on the AFM stage and tests were carried out at room temperature (Figure 1A). AFM tests were performed using a Nanowizard3 AFM (JPK Instruments AG, Germany) equipped with Silicon nitride cantilevers (MLCT, Bruker) and a pyramidal shaped tip. Two cantilevers were used depending on the stiffness of the sample; their nominal spring constant was 0.01 (MLCT-C) and 0.03 (MLCT-D) N/m, respectively. The cantilever spring constant was calibrated using the method described in (*23*) and the deflection sensitivity was determined using the standardized nanomechanical AFM procedure (*24*).

**Figure 1.**
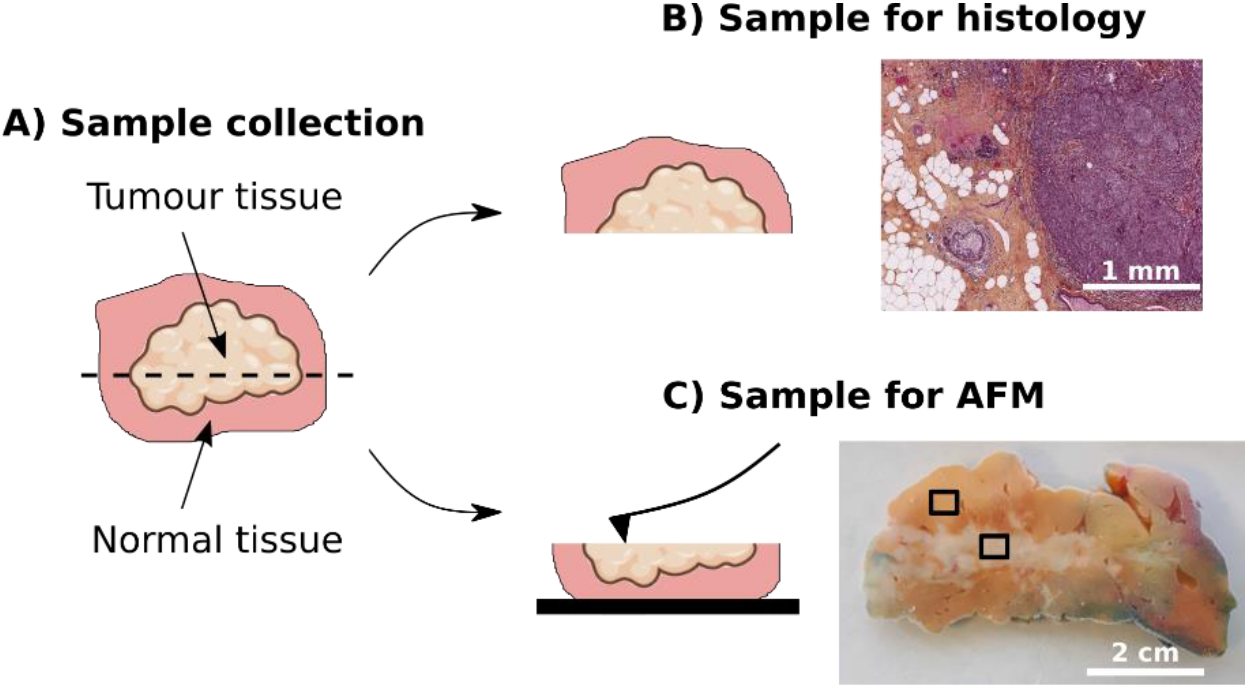
Sample preparation. A) Human samples containing normal and tumour tissue were collected and divided into two samples. B) One sample was used for histology, as part of the routine pathological diagnosis. C) One sample was used for AFM. The photograph shows a breast sample and the black boxes indicate where the AFM measurements were performed.

For each sample, three random regions of at least 1 mm apart were probed over a 100 μm x 100 μm area, with three force maps performed in each region. Since the samples could display high topography, which would have a non-negligible effect on the measurements (*25*), a two-step method was followed (*26*). First, tapping mode topography was used to identify a 15 μm x 15 μm zone of interest with minimal slope (Figure 1B). Second, force-displacement curves were recorded at 3 μm spacing intervals over the zone of interest (25 curves), following a trapezoidal loading profile. The loading segment (extend) was carried out at 2 μm/s until a force of 0.3-0.5 nN was reached. The value of maximal force was adjusted for each sample in order to have an indentation within the range of 300-1500 μm. The maximal force was then held constant for 1 s and the unloading segment (retract) was carried out at 10 μm/s. The range of the force displacement curve was 7 μm to correct the baseline. For each sample, measurement of the mechanical properties was performed within two hours to prevent biochemical changes in the tissue (*10*).

### 2.4 Elastic and viscoelastic analysis

From the raw data (Vertical deflection as a function of piezo height), force-indentation curves were obtained using JPK Data Processing software (v.6.1.163, JPK Instruments AG). The baseline offset and linear fit of the tilt were subtracted from the vertical deflection. If the baseline could not be linearly fitted (e.g. due to a bubble or a particle floating in the medium), the curve was excluded. The contact point, defined as the first crossing of the y axis (vertical deflection), was then identified and the piezo height offset was adjusted. Indentation was calculated by subtracting the cantilever deflection from the piezo height.

Elastic and viscoelastic analyses were then performed from the force-indentation curve and the Indentation-time curve, respectively. For the elastic analysis, the Hertz-Sneddon model for a four-sided pyramid was used, where the <u>**elastic modulus, E**</u>, is defined by equation 1 (*27*).

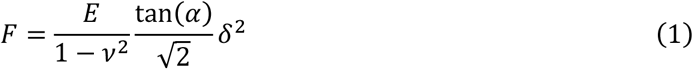

Where F is the force, δ is the indentation, ν is the Poisson’s ratio and α is the face angle of the four-sided pyramid. The elastic modulus represents the stiffness of the material. A Poisson’s ratio of 0.5 was chosen assuming incompressibility of soft biological tissues. The elastic modulus was derived using Equation 1 by fitting the force-indentation curves using E and the contact point as fit parameters. Curves with low quality fit (R^2^<0.98) were discarded. For each sample, values that deviated from the mean by more than three standard deviations were excluded from the analysis.

For the viscoelastic analysis, the standard linear solid (SLS) model was used. The solution for spherical indentation was adapted for indentation with a pyramidal tip (*28*). Briefly, using the elastic-viscoelastic correspondence, equation 1 becomes

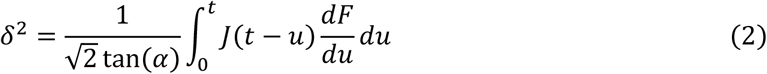

Where u is the dummy variable of integration for time and J(t) is the creep function.

For the SLS model, the creep function is defined as

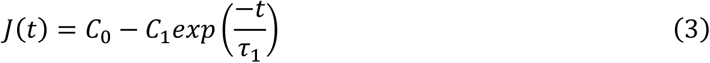

Where *C*_0_, *C*_1_, and *τ*_1_ are model parameters.

For a ramp from zero load at a ramp rate *dP/dt = k*, the solution of equation 2 is

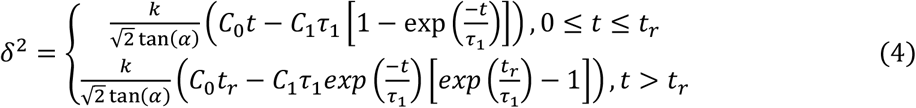

The model parameters were derived using equation 4 by fitting the indentation-time curve on the holding segment. Curves with low quality fit (R^2^<0.95) were discarded.

The instantaneous modulus, *E_ins_*, the equilibrium modulus, *E_eq_*, and the <u>**elastic fraction, *f*,**</u> which quantifies the extent of viscous behaviour of the material (*29, 30*), (with an elastic fraction of 0 representing a purely elastic material, and an elastic fraction of 1 representing a purely viscous material) were then calculated as

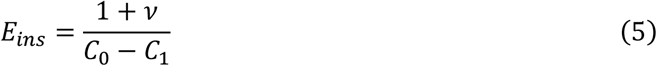

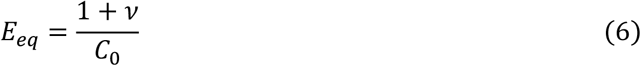

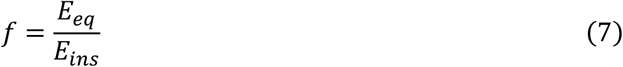

### 2.5 Statistical analysis

Statistical analyses were performed in Python. For each sample, statistical comparison between normal and tumour tissue were made on the total number of indents. Normal distribution was tested using Shapiro test with an alpha risk of 0.05. A Mann-Whitney test was used to compare between normal and tumour tissues the elastic modulus and the elastic fraction with a level of significance of 0.05.

## 3 Results

### 3.1 Mechanical properties and structure of breast tumours

Breast tumour samples were composed of invasive breast carcinoma of no special type (NST) (n=3) and invasive lobular carcinomas (ILC) (n=3) (Figure 3). Mean and median values of elastic modulus and elastic fraction for each sample are reported in Table 1. Elastic modulus of tumour tissue significantly increased compared to its normal counterpart in 5 out of 6 samples, with mean values of 1.4 ± 0.91 kPa in control and 3.3 ± 3.4 kPa in tumour tissue for NST, and 1.2 ± 0.29 kPa in control and 2.8 ± 1.5 kPa in tumour tissue for ILC (Figure 3A, Table 1). No particular trend was observed in the elastic fraction between normal and tumour tissue, with mean values of 0.53 ± 0.069 in control and 0.53 ± 0.047 in tumour tissue (Figure 3B, Table 1). Only one sample displayed a significant increase in the elastic fraction in tumour tissue. NST samples displayed a well-defined tumour mass within the adipose tissue (Figure 3C-D), while ILC samples displayed a more diffuse tumour mass (Figure 3E-F). Both types exhibited an extensive fibrotic stroma.

**Figure 2.**
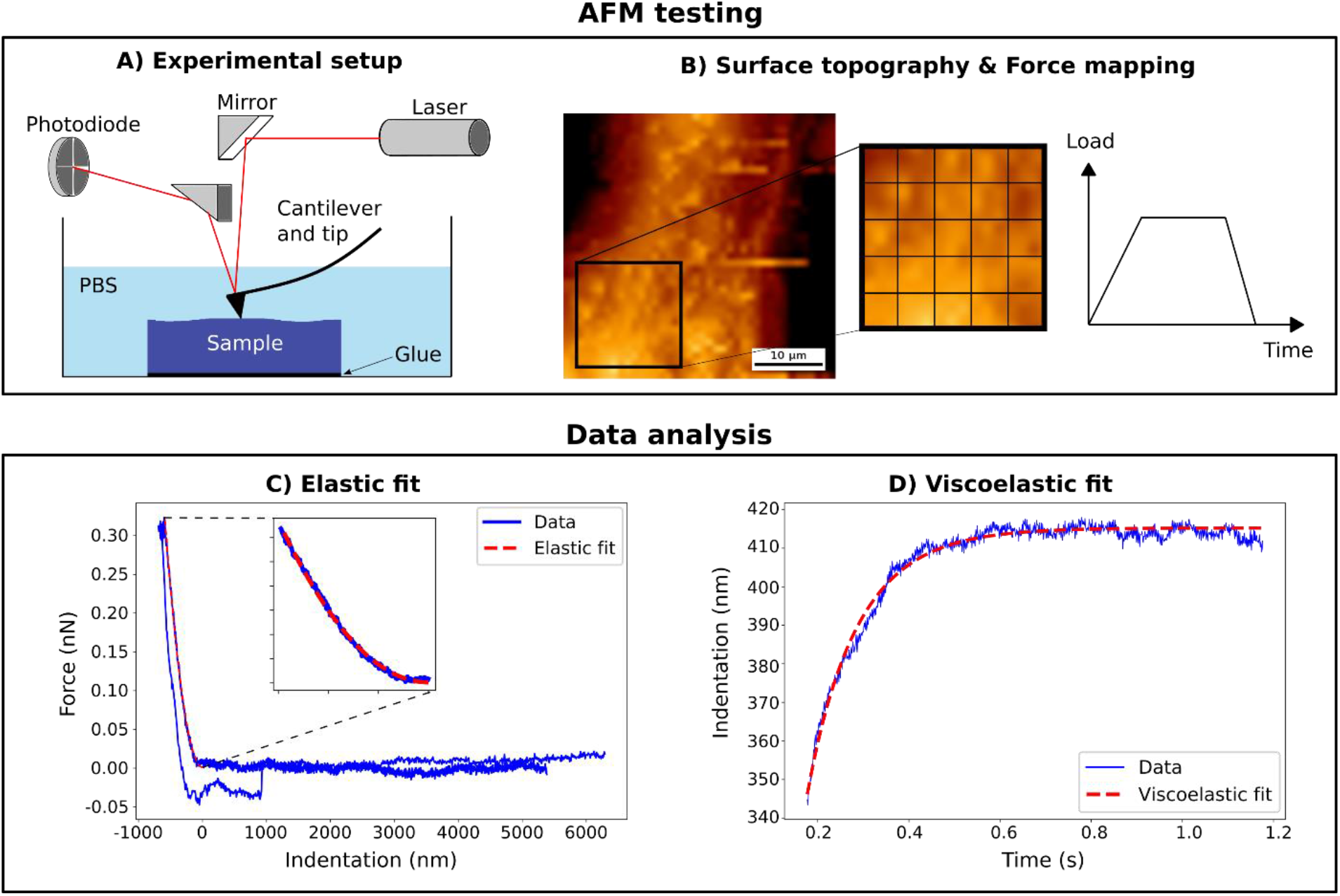
Method overview for the mechanical characterisation. A) Schematic of the AFM setup. The sample was glued on a petri dish, immersed in PBS and placed on the AFM scanner. B) The AFM taping mode topography was first used to get the surface profile and identify a region of interest (black box). The force mapping mode was then used in this region of interest with a trapezoidal loading profile. C) Typical force-indentation curve obtained from AFM measurements (blue). Elastic modulus was obtained by fitting the extend segment using Hertz model (red). D) Typical Indentation-time curve on the holding segment (blue). Elastic fraction was obtained by fitting the holding segment using Maxwell model (red).

**Figure 3.**
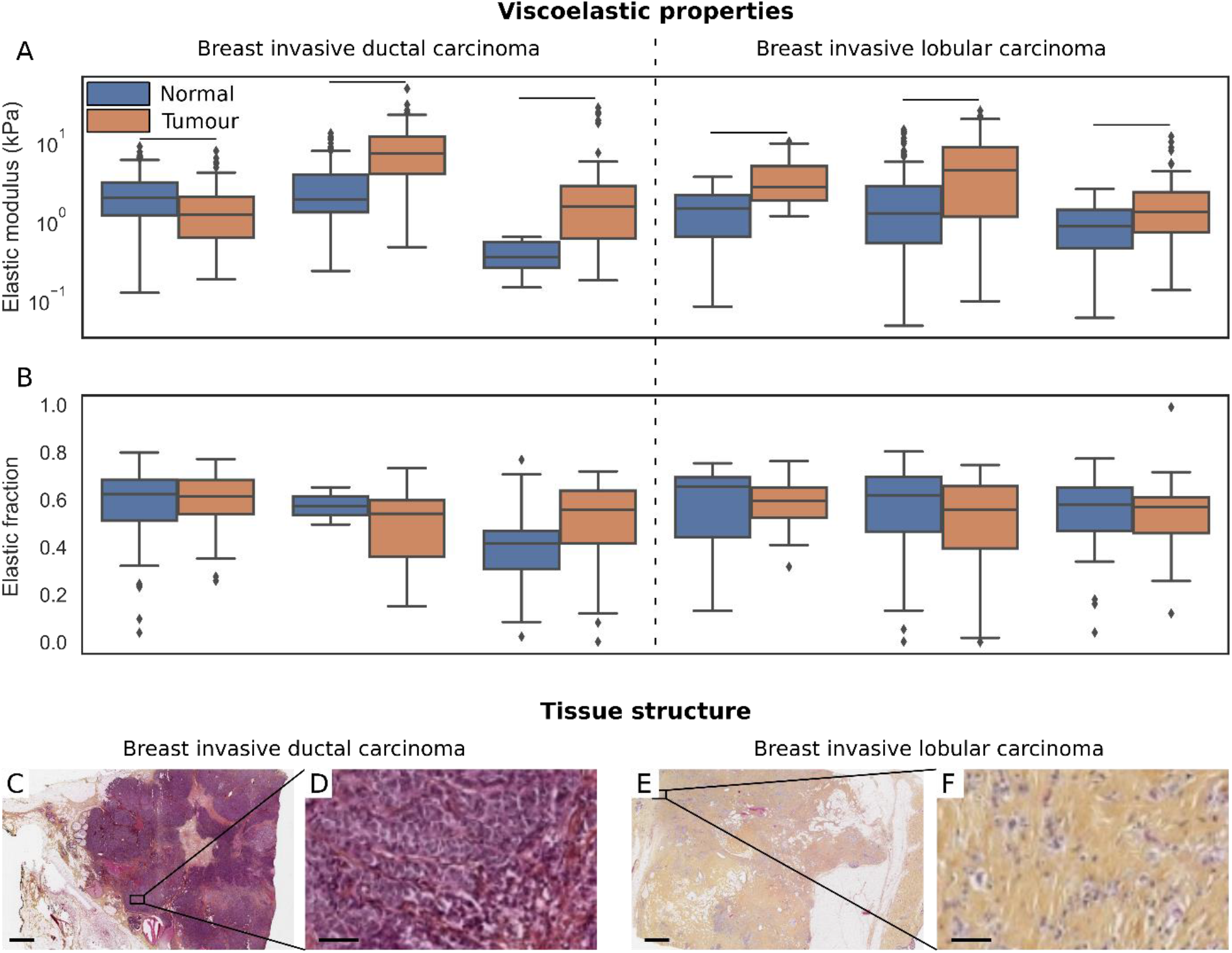
Breast tumour samples, which were composed of dense fibrous tissue, displayed an increased elastic modulus compared to their normal counterpart. Viscoelastic properties and histopathological examination of breast tumour tissue. A) Elastic modulus and B) elastic fraction of normal (blue) and tumour (orange) tissue for two types of breast cancers and different patients. C-J) Hematoxylin eosin saffron stained sections of breast tumour tissue of each type. Horizontal bar indicates significant difference between normal and tumour samples. Scale bars: 2 mm (C, E) and 60 μm (D, F).

**Table 1:** Elastic modulus and elastic fraction of normal and tumour tissue for each breast sample. Results are indicated as mean ± standard deviation (std) and median. The p values show differences between normal and tumour tissue for each sample. NST: No Special type; ILC: Invasive Lobular Carcinoma.

| Breast |  |  |  |  |  |  |
| --- | --- | --- | --- | --- | --- | --- |
| Sample # | Type | Group | Elastic modulus (kPa) |  | Elastic fraction |  |
| | | | Mean ± Std | Median | Mean ± Std | Median |
| 1 | NST | Normal | 2.4 ± 1.7 | 2.0 | 0.58 ± 0.16 | 0.62 |
| | | Tumour | 1.5 ± 1.3 | 1.2 | 0.60 ± 0.10 | 0.61 |
|  |  |  | **p < 0.001** |  | p = 0.78 |  |
| 2 | NST | Normal | 3.2 ± 3.0 | 1.9 | 0.57 ± 0.11 | 0.57 |
| | | Tumour | 8.5 ± 6.7 | 7.2 | 0.48 ± 0.16 | 0.54 |
|  |  |  | **p < 0.001** |  | p = 0.42 |  |
| 3 | NST | Normal | 0.37 ± 0.16 | 0.35 | 0.40 ± 0.16 | 0.41 |
| | | Tumour | 2.7 ± 4.4 | 1.5 | 0.50 ± 0.17 | 0.56 |
|  |  |  | **p < 0.001** |  | **p < 0.001** |  |
| 4 | ILC | Normal | 1.4 ± 0.90 | 1.4 | 0.56 ± 0.18 | 0.66 |
| | | Tumour | 3.7 ± 2.8 | 2.7 | 0.58 ± 0.14 | 0.60 |
|  |  |  | **p < 0.001** |  | p = 0.88 |  |
| 5 | ILC | Normal | 2.5 ± 3.2 | 1.2 | 0.56 ± 0.19 | 0.62 |
| | | Tumour | 5.9 ± 5.6 | 4.4 | 0.51 ± 0.19 | 0.60 |
|  |  |  | **p < 0.001** |  | p = 0.068 |  |
| 6 | ILC | Normal | 0.93 ± 0.64 | 0.85 | 0.53 ± 0.17 | 0.58 |
| | | Tumour | 1.9 ± 2.1 | 1.3 | 0.54 ± 0.20 | 0.60 |
|  |  |  | **p < 0.001** |  | p = 0.70 |  |

### 3.2 Mechanical properties and structure of **kidney tumours**

We obtained 5 kidney tumour samples composed of clear cell renal cell carcinomas (RCC) (n=3), papillary RCC (n=1) and a mix between clear cell RCC and sarcomatoid (n=1) (Figure 4). Mean and median values of elastic modulus and elastic fraction for each sample are reported in Table 2. Elastic modulus of tumour tissue significantly decreased compared to its normal counterpart for all clear cell RCC samples, with mean values of 3.2 ± 2.7 kPa in control and 1.7 ± 1.6 kPa in tumour tissue (Figure 4A). The elastic fraction was similar between normal (mean value: 0.48 ± 0.14) and tumour (mean value: 0.52 ± 0.021) tissue, with only one sample showing a significant decrease in tumour tissue. Tumour tissue exhibited a high cellular density with little fibrotic tissue compared to normal tissue (Figure 4C-D). Papillary RCC sample displayed a significant increase in the elastic modulus in tumour tissue compared to normal tissue, with a high heterogeneity in tumour tissue (Figure 4A, Table 2). The mean elastic modulus was 67 ± 138 kPa in tumour and 3.0 ± 4.5 kPa in control tissue. No significant difference was observed in the elastic fraction between normal and tumour tissue. Cancer cells were packed in a papillary architecture and surrounded by fibrotic tissue (Figure 4E-F). The sample with a mix of clear cell and sarcomatoid RCC showed a high heterogeneity in the elastic properties in tumour tissue, with a mean elastic modulus of 15 ± 36 kPa (Figure 4A, Table 2). No significant difference was observed in the elastic modulus and elastic fraction between normal and tumour tissue (Figure 4B, Table 2). This sample exhibited a high cellular density with little fibrotic tissue in the clear cell carcinoma area, and elongated cells surrounded by a fibrotic stroma in the sarcomatoid carcinoma area (Figure EG-H).

**Figure 4.**
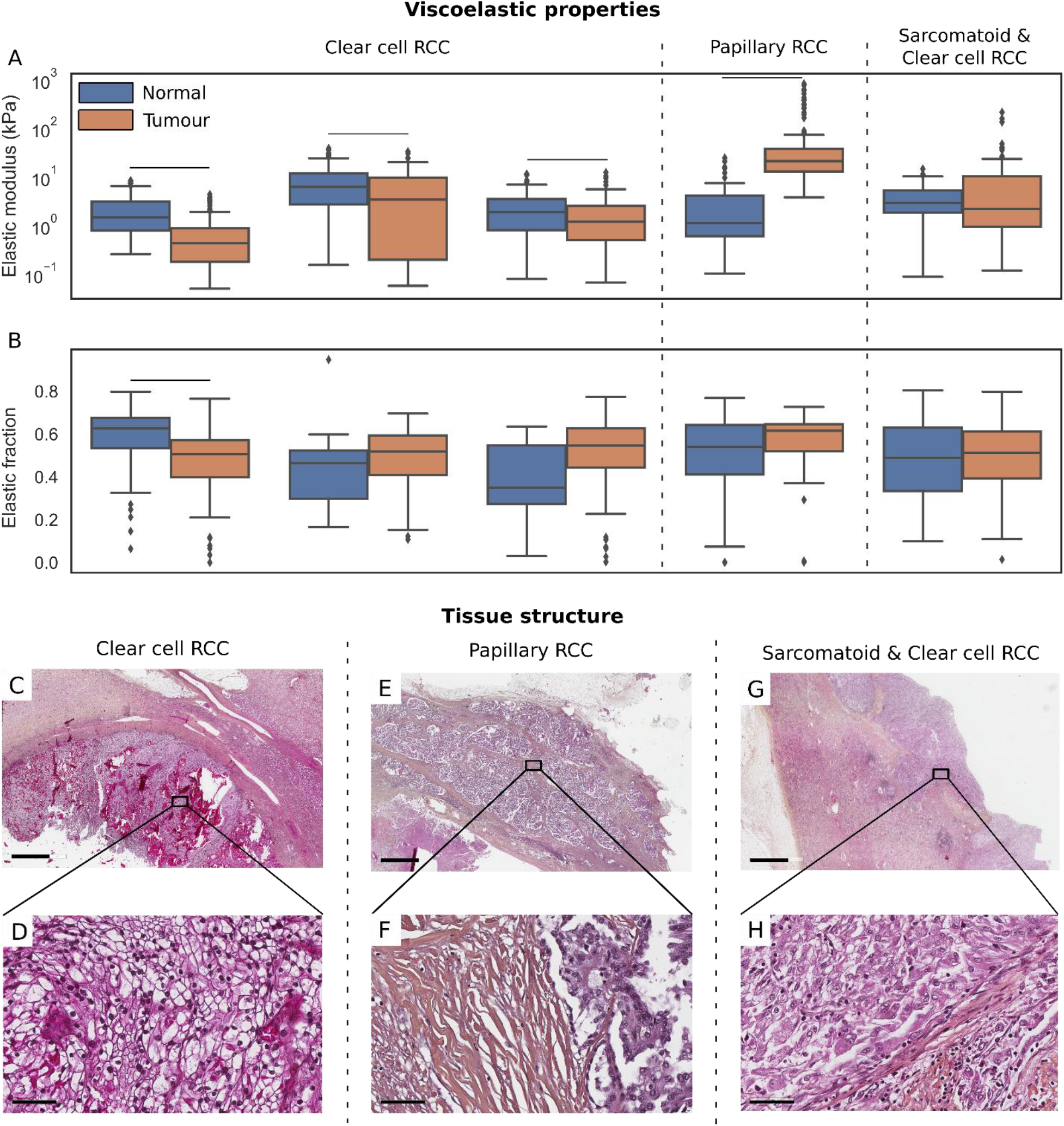
Elastic properties of kidney tumour samples were differently affected depending on their histological type. Viscoelastic properties and histopathological examination of renal tumour tissue. A) Elastic modulus and B) elastic fraction of normal (blue) and tumour (orange) tissue for three types of kidney cancers and different patients. C-J) Hematoxylin eosin saffron stained sections of kidney tumour tissue of each type. Horizontal bar indicates significant difference between normal and tumour samples. RCC: Renal Cell Carcinoma. Scale bars: 2 mm (C, E) and 60 μm (D, F).

**Table 2:** Elastic modulus and elastic fraction of normal and tumour tissue for each kidney sample. Results are indicated as mean ± standard deviation (std) and median. The p values show differences between normal and tumour tissue for each sample. CC: Clear Cell renal cell carcinoma; P: Papillary carcinoma; CC-S: Clear Cell and Sarcomatoid carcinoma.

| Kidney |  |  |  |  |  |  |
| --- | --- | --- | --- | --- | --- | --- |
| Sample # | Type | Group | Elastic modulus (kPa) |  | Elastic fraction |  |
| | | | Mean ± Std | Median | Mean ± Std | Median |
| 1 | CC | Normal | 2.2 ± 1.9 | 1.5 | 0.59 ± 0.14 | 0.63 |
| | | Tumour | 0.75 ± 0.85 | 0.44 | 0.47 ± 0.16 | 0.51 |
|  |  |  | **p < 0.001** |  | **p < 0.001** |  |
| 2 | CC | Normal | 8.9 ± 8.4 | 6.3 | 0.44 ± 0.22 | 0.46 |
| | | Tumour | 6.7 ± 8.8 | 3.5 | 0.47 ± 0.17 | 0.52 |
|  |  |  | **p = 0.0073** |  | p = 0.33 |  |
| 3 | CC | Normal | 2.8 ± 3.0 | 1.9 | 0.37 ± 0.21 | 0.35 |
| | | Tumour | 1.9 ± 2.1 | 1.2 | 0.51 ± 0.17 | 0.55 |
|  |  |  | **p = 0.028** |  | p = 0.055 |  |
| 4 | P | Normal | 3.0 ± 4.5 | 1.1 | 0.49 ± 0.20 | 0.54 |
| | | Tumour | 67 ± 139 | 21 | 0.55 ± 0.17 | 0.62 |
|  |  |  | **p < 0.001** |  | p = 0.17 |  |
| 5 | CC-S | Normal | 3.9 ± 3.2 | 2.9 | 0.47 ± 0.19 | 0.49 |
| | | Tumour | 15 ± 36 | 2.2 | 0.49 ± 0.16 | 0.51 |
|  |  |  | **p = 0.79** |  | p = 0.77 |  |

### 3.3 Mechanical properties and structure of **thyroid tumours**

Thyroid tumour samples were composed of papillary (n=1), anaplastic (n=1) and poorly differentiated (n=1) carcinomas, with one additional sample exhibiting papillary and anaplastic types. Mean and median values of elastic modulus and elastic fraction for each sample are reported in Table 3. The papillary carcinoma and anaplastic samples displayed a significantly increased elastic modulus in tumour tissue compared to their normal counterpart, with mean values of 0.54 ± 0.28 kPa in control and 1.7 ± 1.7 kPa in tumour tissue for papillary carcinoma, and 1.3 ± 1.0 kPa in control and 2.4 ± 2.2 kPa in tumour tissue for anaplastic carcinoma (Figure 5A, Table 3). The papillary carcinoma sample also displayed a decrease in the elastic fraction in tumour tissue, while no difference was observed for the anaplastic carcinoma sample. In both samples, cancer cells were surrounded by an extensive fibrotic stroma (Figure 5C-F). The papillary/anaplastic and poorly differentiated samples displayed a significantly decreased elastic modulus in tumour tissue compared to its normal counterpart, with mean values of 1.4 ± 1.6 kPa in control and 0.070 ± 0.056 kPa in tumour tissue for papillary/anaplastic carcinoma, and 2.9 ± 2.5 kPa in control and 0.78 ± 0.54 kPa in tumour tissue for poorly differentiated carcinoma (Figure 5A, Table 3). Both samples exhibited a high cellular density with little fibrotic tissue (Figure 5G-J).

**Figure 5.**
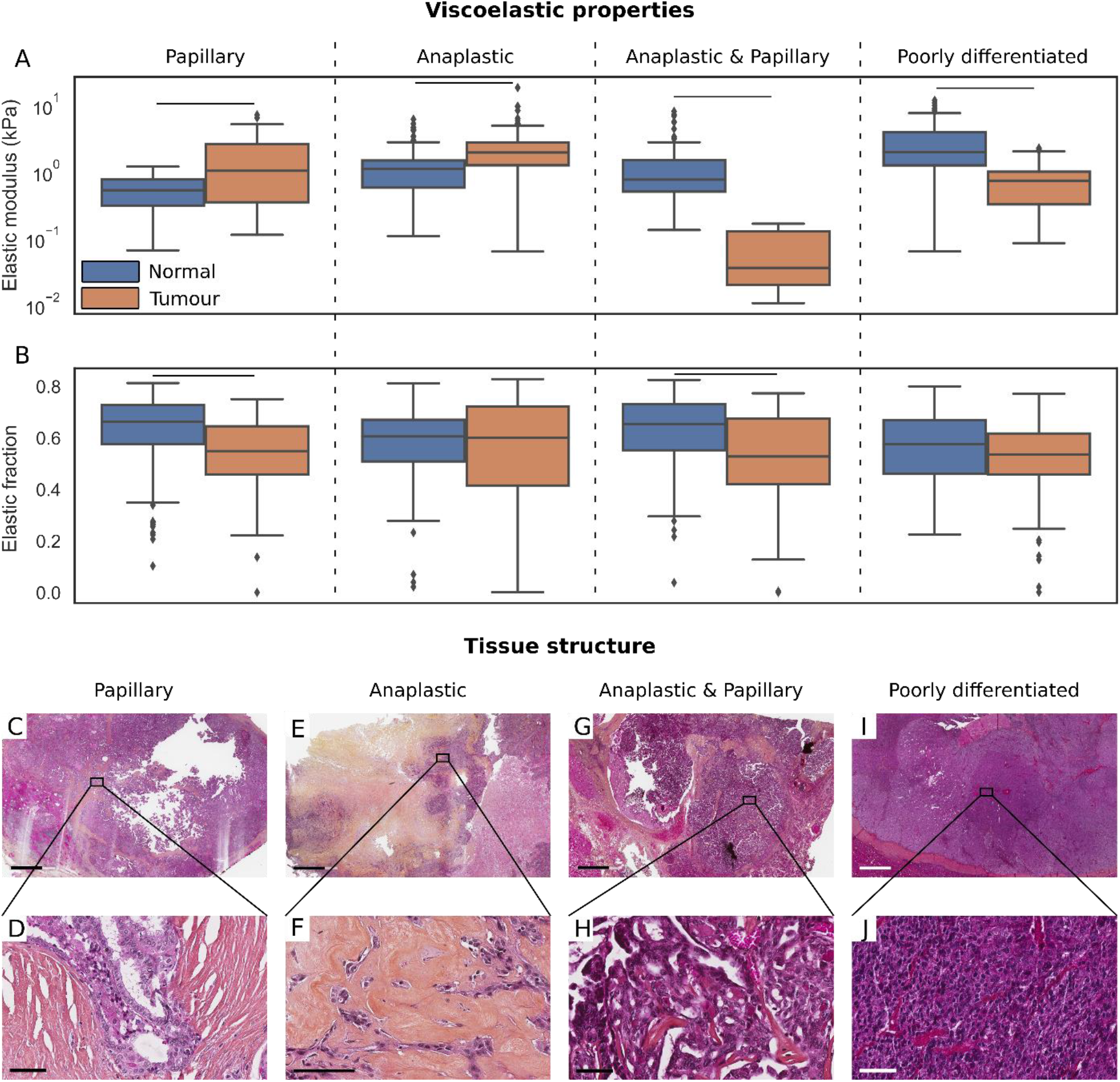
Elastic properties of thyroid tumour samples were differently affected depending on their histological type, while the elastic fraction tended to decrease in most tumour samples. Viscoelastic properties and histopathological examination of thyroid tumour tissue. A) Elastic modulus and B) elastic fraction of normal (blue) and tumour (orange) tissue for different types of thyroid cancers. C-J) Hematoxylin eosin saffron stained sections of thyroid tumour tissue. Horizontal bar indicates significant difference between normal and tumour samples. Scale bars: 2 mm (top row) and 60 μm (bottom row).

**Table 3:**
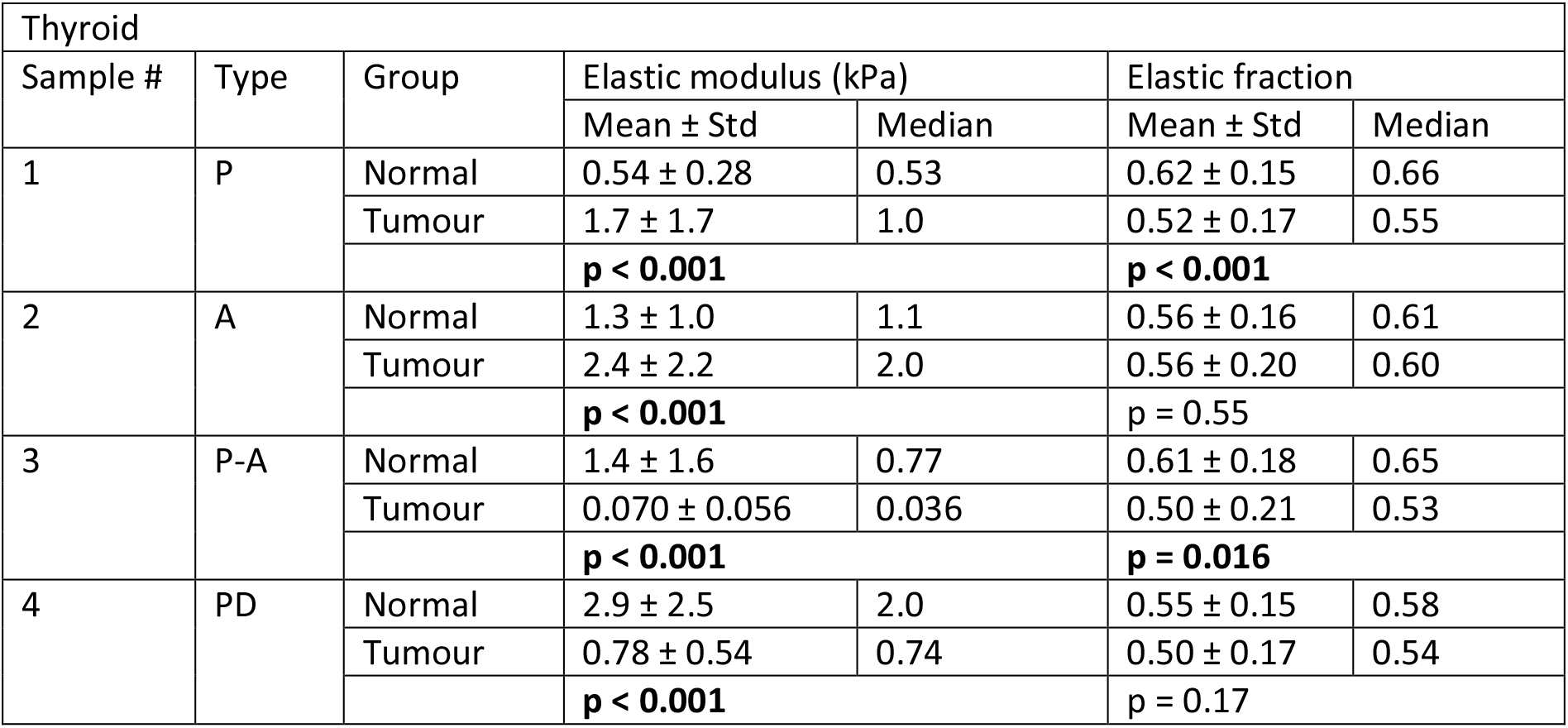
Elastic modulus and elastic fraction of normal and tumour tissue for each thyroid sample. Results are indicated as mean ± standard deviation (std) and median. The p values show differences between normal and tumour tissue for each sample. P: Papillary carcinoma; A: Anaplastic carcinoma; P-A: Papillary and Anaplastic carcinoma; PD: Poorly Differentiated carcinoma.

## 4 Discussion

In the present study, we showed that elastic and viscoelastic properties of tumour tissue were differentially affected depending on the organ and on the structure and composition of the tissue. The first hypothesis, that the elastic properties of the tumours were differentially affected depending on their type, was corroborated. Clear cell renal carcinoma and poorly differentiated thyroid carcinoma displayed a decrease in the elastic modulus compared to their normal counterpart, while breast tumours, papillary renal carcinoma and fibrotic thyroid tumours displayed an increase in the elastic modulus. The second hypothesis, that tumour tissue exhibits a change in viscous properties compared to normal tissue, was partially corroborated. While the elastic fraction of thyroid tumour tissue tended to be lower compared to their normal counterpart, indicating an increase in the viscosity, no consistent difference was observed for breast and kidney tissue. These findings suggest a unique mechanical profile associated with each subtype of cancer.

The present study emphasizes the potential of AFM to characterise tissue mechanical properties of different types of tumours. We developed a method to identify both elastic and viscoelastic properties, providing new valuable information for the development of biomaterials mimicking the mechanical properties of the tumour microenvironment. Indentation tests using AFM were performed on intact samples, without any processing to make the surface smooth or levelling the sample. The surface tomography obtained prior to the tests ensured that the slope and the surface irregularities on the zone of interest were kept minimal. The elastic modulus of breast tissues obtained using this protocol was in agreement with other studies using AFM (*7, 10*). For example, our mean values of elastic modulus for normal (1.4 kPa) and tumour (3.3 kPa) breast tissues were consistent with the study of Plodinec *et al*, who obtained peak stiffness values of 1.13 to 1.83 kPa for healthy tissue, and 1.54 to 9.62 kPa for malignant breast tumour tissue (*7*). For the elastic fraction, the values for tumour breast tissue obtained in our study ranged between 0.48 and 0.60 and were higher than the values obtained in Qiu *et al*, which ranged between 0.15 and 0.38. However, in their study, Qiu *et al* characterized the viscoelastic properties of tumour tissue obtained after subcutaneously injecting breast cancer cells in a murine model (*31*). Therefore, the differences in the microenvironment between breast and subcutaneous tissues can explain the different mechanical properties between both tumour tissues. To our knowledge, mechanical properties of thyroid and kidney tumour tissues had not been characterised using AFM. Although the elastic modulus values found in our study were lower than the values obtained using compression testing (*14*), shear wave elastography (SWE) (*32*), or palpation device technique (*13*), which can be explained by the different measurement scales, the comparison between healthy and tumour tissue was consistent with the literature. Lee *et al* found that clear cell renal cell carcinoma tissues displayed a decrease in their elastic modulus compared to benign tissue, with mean values of 13.0 ± 8.2 kPa and 19.2 ± 10.8 kPa, respectively (*13*). For thyroid tissue, measurements at the macroscopic scale revealed a higher elastic modulus of papillary tumour tissue compared to normal tissue (*14, 32*), which is consistent with our findings.

The results of this study demonstrate that the mechanical properties of tumour tissues are differentially affected depending on their type and subtype, and that the stiffness of tumour tissue is directly associated with ECM remodelling. Breast tumour tissues displayed a dense fibrotic stroma, resulting in a significant increase in the elastic modulus, which was twice to three times higher than the elastic modulus of normal breast samples. This increase in the elastic modulus seemed to be stronger for NST samples, which were characterised by a well-defined and dense tumour mass, compared to ILC samples. Similarly, papillary renal carcinoma, papillary and anaplastic thyroid tumours exhibited a fibrotic stroma, associated with an increase in the elastic modulus compared to their normal counterpart. Tumour fibrosis is a well-known feature associated with cancer progression and stiffening of the stroma (*33*). Indirectly, we have previously observed that phenomenon in our POUMOS cohort with periostin, a marker of ECM reaction that was associated with survival in bone metastatic adenocarcinoma lung cancer patients (*34*). The disruption of the balance between ECM synthesis and enzymatic degradation in favour of ECM synthesis has been primarily attributed to the inability of matrix metalloproteinases (MMPs) to digest collagen and alteration of the mode of collagen cross-linking (*35*). By contrast, clear cell renal carcinoma and poorly differentiated thyroid carcinoma displayed a high cellular density, which was associated with a decrease in the elastic modulus by at least 50%. Further investigation at the tissue and molecular level will be necessary to elucidate the mechanisms behind this imbalance of ECM remodelling in favour of enzymatic degradation. Altogether, these findings suggest specific processes by which cancer cells alter the structure and composition of tumour tissue, resulting in a different effect on the mechanical properties of the tissue. The type of changes associated with each cancer likely affects cancer progression and metastasis formation.

Several studies suggest a link between the mechanical properties of breast tumour and their metastatic potential and aggressiveness. In particular, it has been shown using a mouse model of breast cancer that tissue stiffening can promote bone metastasis (*36*). Similarly, several studies indicated a positive correlation between breast cancer aggressiveness and tissue stiffening at the macroscale (*9*) and tissue scale (*37*). The relationships between tumour stiffness and its metastatic potential and aggression have not been investigated for thyroid and kidney cancer. Our findings suggest different conclusions than for breast tissue. Indeed, we found that poorly differentiated thyroid and anaplastic carcinomas, respectively, which are very aggressive and frequently metastasise to bone and lung (*38, 39*), follow different trends, with either a reduced or increased elastic modulus compared to their normal counterpart, respectively. Moreover, we showed that clear cell renal carcinomas, which have the highest metastatic potential (*40*), display a reduced elastic modulus, while papillary carcinoma displays an increased elastic modulus. Therefore, the relationships between tissue stiffness and cancer aggressiveness and metastatic potential merit further investigation for thyroid and kidney cancer.

Our study also provides new knowledge on the viscoelastic properties of tumour tissue. Although the viscous properties of the ECM are known to play a role for cell activities, very few studies have characterised the viscoelastic properties of tumour tissue. In their *in vivo* study investigating the viscoelastic shear properties of pathological breast tissue using magnetic resonance elastography (MRE), Sinkus *et al* showed that the shear modulus was a better parameter than the viscosity at differentiating between benign and malignant lesions (*41*). Moreover, they found two clusters of breast cancer, one of which was as viscous as the surrounding tissue. Conversely, Garteiser *et al* found that the loss modulus of liver tumours measured using MRE, which represents the viscous behaviour of the tissue, was a better discriminator between benign and malignant tumours than the storage modulus, which represents the elastic behaviour (*42*). These findings demonstrate that the type of viscoelastic change allowing the discrimination between benign and malignant tumours depend on the tissue. Similarly, in our study at the tissue level, we found discrepancies in the viscoelastic changes between normal and tumour tissues depending on the type of cancer. While breast and kidney tumours did not exhibit any consistent difference in their elastic fraction compared to their surrounding normal tissue, thyroid tumour tissues tended to be more viscous than their surrounding normal tissue. Therefore, viscosity could be a discriminator between tumour and normal thyroid tissue, while elasticity could be a discriminator between the subtypes of cancer.

The current study presents several limitations. First, the robustness of the results is limited by the small sample size, which is mainly due to the difficulty in obtaining tissue from specific subtypes of cancer. In particular, some cancer types are rare (sarcomatoid renal carcinoma, anaplastic and poorly differentiated thyroid carcinomas), and only one sample for each of these types could be obtained. Second, due to logistic reasons, the samples were frozen before their mechanical characterisation. There is no consensus on the literature as to whether freezing affects the mechanical properties of biological soft tissues. Indeed, some studies showed that there is not any significant difference in the stiffness between fresh and frozen tissues (*43–46*), while other studies found an increase (*47*) or a decrease (*48*) in the stiffness of frozen tissue compared to fresh tissue. The elastic modulus of breast tissue obtained in our study were in agreement with the values obtained on fresh tissue (*7, 10*), suggesting that freezing did not significantly alter the mechanical properties of these tumour tissues. Moreover, we proposed a comparative study with the same conservation mode for all the tissues. Third, since the current study focused on the mechanical properties of tumour tissues, assessment of tissue structure using histology has not been performed on the same samples. Future studies will aim to spatially correlate tissue structure and mechanical properties by performing AFM and histology on the same samples, following cryosectioning for example (*49*). Moreover, an investigation at the ultra-structural and molecular scales needs to be done to understand the mechanisms of how the mechanical properties of each type of tumour are differentially affected.

From a clinical perspective, our findings on the mechanical properties of tumour tissue have the potential to significantly advance the field of cancer research and have implications for diagnosis purposes. A recent study developing hydrogels mimicking the heterogeneous mechanical properties of breast tumour showed how stiffness regulates pro-metastatic functions of breast cancer cells, both *in vitro* and *in vivo* (*50*). Our findings on the mechanical properties of breast, kidney, and thyroid tumour tissue will allow the development of similar biomaterials, in order to elucidate the role of stiffness and viscosity for metastatic spread depending on cancer subtype. With the progress of *in vivo* imaging techniques to characterise the viscoelastic properties of soft tissues with a higher spatial resolution, namely SWE (*51*) and microscopic Magnetic Resonance Elastography (MRE) (*52–54*), the mechanical properties of the tumour could then be used as an *in vivo* predictive marker of bone metastasis formation.In conclusion, this study elucidated the modification of the viscoelastic properties of tumour tissue from different types of cancer. While the elastic modulus of breast tumour tissue increased compared to the normal surrounding tissue, various effects were observed for kidney and thyroid tissue depending on the type, structure and composition of the tumour. The elastic fraction of tumour tissue decreased only for the thyroid, suggested a role of viscosity for thyroid cancer progression. These findings on different types of cancer form a first basis to understand how mechanical alterations are linked to aggressiveness and metastatic potential of cancer. Identification of mechanical parameters predictive of cancer progression will have implications for diagnosis purposes.

## 5. Acknowledgments

We thank Corinne Perrin and Karine Castellano from Hospices Civils de Lyon for specimen collection, Sara Calattini and Mélanie Roche from Cancer Institute Research Platform of Hospices Civils de Lyon (IC-HCL) for coordinating the clinical study. We also acknowledge the contribution of SFR Biosciences (UAR3444/CNRS, US8/Inserm, ENS de Lyon, UCBL) facilities and the staff of Platim, especially Simone Bovio for his help with AFM. This study has received funding from the European Union’s Horizon 2020 research and innovation programme under the Marie Skłodowska-Curie grant agreement No. 895139 and MSDAvenir research grant (MEKANOS project).

